# The wake- and sleep-modulating neurons of the lateral hypothalamic area demonstrate a differential pattern of degeneration in Alzheimer’s disease

**DOI:** 10.1101/2024.03.06.583765

**Authors:** Abhijit Satpati, Felipe L. Pereira, Alexander V. Soloviev, Mihovil Mladinov, Eva Larsen, Song Hua Li, Chia-Ling Tu, Renata E. P. Leite, Claudia K. Suemoto, Roberta D. Rodriguez, Vitor R. Paes, Christine Walsh, Salvatore Spina, William W. Seeley, Carlos A. Pasqualucci, Wilson Jacob Filho, Wenhan Chang, Thomas C. Neylan, Lea T. Grinberg

**Affiliations:** Memory and Aging Center, University of California San Francisco, USA; Skeletal Biology and Biomechanics Core, University of California San Francisco, USA; Physiopathology in Aging Laboratory (LIM-22), Department of Internal Medicine, University of Sao Paulo Medical School, Sao Paulo, Brazil; Department of Pathology, University of Sao Paulo Medical School, Sao Paulo, Brazil; Division of Geriatrics, Department of Internal Medicine, University of Sao Paulo Medical School, Sao Paulo, Brazil; University of São Paulo Medical School, São Paulo, Brazil; Department of Psychiatry, University of California San Francisco, San Francisco, California, USA

**Keywords:** Digital multiplexed gene expression, Melanin-concentrating hormone, Orexin/Hypocretin, RNA expression level of LHA, Unbiased stereology

## Abstract

**Background:** Sleep-wake dysfunction is an early and common event in Alzheimer’s disease (AD). The lateral hypothalamic area (LHA) regulates the sleep and wake cycle through wake-promoting orexinergic neurons (Orx^N^) and sleep-promoting melanin-concentrating hormone or MCHergic neurons (MCH^N^). These neurons share close anatomical proximity with functional reciprocity. This study investigated LHA Orx^N^ and MCH^N^ loss patterns in AD individuals. Understanding the degeneration pattern of these neurons will be instrumental in designing potential therapeutics to slow down the disease progression and remediate the sleep-wake dysfunction in AD.

**Methods:** Postmortem human brain tissue from donors with AD (across progressive stages) and controls were examined using unbiased stereology. Formalin-fixed, celloidin-embedded hypothalamic sections were stained with Orx-A/MCH, p-tau (CP13), and counterstained with gallocyanin. Orx or MCH-positive neurons with or without CP13 inclusions and gallocyanin-stained neurons were considered for stereology counting. Additionally, we extracted RNA from the LHA using conventional techniques. We used customized Neuropathology and Glia nCounter^®^ (Nanostring) panels to study gene expression. Wald statistical test was used to compare the groups, and the genes were considered differentially expressed when the p-value was <.05.

**Results:** We observed a progressive decline in Orx^N^ alongside a relative preservation of MCH^N^. Orx^N^ decreased by 58% (p=.03) by Braak stages (BB) 1-2 and further declined to 81% (p=.03) by BB 5-6. Conversely, MCH^N^ demonstrated a non-statistical significant decline (27%, p=.1088) by BB 6. We observed a progressive increase in differentially expressed genes (DEGs), starting with glial profile changes in BB2. While Orx^N^ loss was observed, Orx-related genes showed upregulation in BB 3-4 compared to BB 0-1. GO and KEGG terms related to neuroinflammatory pathways were mainly enriched.

**Conclusions:** To date, Orx^N^ loss in the LHA represents the first neuronal population to die preceding the loss of LC neurons. Conversely, MCHN shows resilience to AD p-tau accumulation across Braak stages. The initial loss of Orx^N^ correlates with specific neuroinflammation, glial profile changes, and overexpression of HCRT, possibly due to hyperexcitation following compensation mechanisms. Interventions preventing Orx^N^ loss and inhibiting p-tau accumulation in the LHA could prevent neuronal loss in AD and, perhaps, the progression of the disease.

## Introduction

The lateral hypothalamic area (LHA), delimited in the human brain by the dorsal fornix, mamillotegmental and mamillotalamic tracts, and the third ventricle, is a pivotal brain region regulating sleep and wakefulness, and feeding behaviors. The LHA has gained increased attention following the discovery of two distinct neuronal populations: Orexin (Orx)/Hypocretin (Hcrt) neurons in 1998 ^1,2^ and melanin-concentrating hormone (MCH) neurons in the 1980s ^3^. Orx and MCH, neuronal populations mostly unique to the LHA, are essential for coordinating sleep with other vital functions and integrating these functions into a 24-hour circadian rhythm, making the LHA a significant focus of neuroscientific research.

The neuropeptide Orexin (Orx)/Hypocretins (Hcrt) is produced by 50,000 - 80,000 orexinergic neurons (Orx^N^) maintains wakefulness, and stimulates appetite ^4–7^. Orx regulates glutamate and ACh release in the frontal and prefrontal cortex and mediates the presynaptic component and peptide control of fast-spiking interneurons ^8,9^. Orx^N^ also offers neuroprotective benefits through its anti-inflammatory actions. Loss of Orx^N^ underlies narcolepsy ^10,11^. MCH neurons (MCH^N^) play a critical role in maintaining energy balance ^12^, and REM sleep ^13^, demonstrating their physiological importance across species from fish to mammals. However, despite their critical roles, MCH^N^ remains largely uncharacterized in humans.

Alzheimer’s disease (AD) neuropathology is the most common neuropathological basis for dementia, affecting millions worldwide. While short-term memory loss is the most recognized early symptom linked to the progression of AD neuropathological changes, a considerable number of individuals also experience sleep-wake disturbances and other neuropsychological symptoms, which often precede memory impairments. Historically, due to limited understanding of the neuropathological foundations of these symptoms, they were primarily considered risk factors for AD-type dementia. However, recent clinicopathological and imaging studies indicate that AD pathology, particularly AD-tau pathology, may actually underpin these neuropsychological symptoms, suggesting that they are part of the AD spectrum itself. For example, in a study involving over 1,000 individuals from a population-based clinicopathological cohort, those with AD Braak stage I/II (i.e., very mild cortical changes and subcortical tau changes) exhibited more than twice the likelihood of experiencing appetite disturbances and nighttime behaviors compared to individuals at Braak stage 0 (i.e., mild subcortical tau changes only) ^14^. Previous research by our group and others has shown that Orx^N^ accumulates AD-tau from Braak 0. Stereological assessment on individuals at later Braak stages and healthy controls demonstrated a 40-70% reduction in the Orx^N^ population ^15,16^. Furthermore, the loss of Orx^N^ was associated with increased total sleep time, more frequent wakefulness after sleep onset, a slight reduction in time spent in the N3 sleep stage, and a decrease in the percentage of time spent in REM sleep ^17^. Collectively, these findings suggest that Orx^N^ dysfunction plays a role in the pathogenesis of AD from its earliest stages. However, it remains unclear how early Orx^N^ begins to degenerate in AD and the specific molecular mechanisms underlying AD-tau associated Orx^N^ loss - a significant knowledge gap, especially given that paradoxically, moderate to severe AD patients with sleep impairments and cognitive decline often exhibit higher CSF and plasma orexin levels. This elevated CSF orexin correlates positively with CSF Aβ42, total tau (t-tau), and phosphorylated tau (p-tau) levels ^18–22^ and could suggest a compensatory mechanism designed to mitigate the effects of Orx^N^ loss, the underlying mechanisms of which are also not well understood.

Despite extensive research on vertebrates, very little is known about MCH^N^, not only in AD but also in the human brain in general. While animal studies have shed valuable light on the orexinergic and MCH systems, in AD, the LHA primarily accumulates AD-tau rather than beta-amyloid. Unfortunately, no animal model available accurately represents AD-tau accumulation, posing significant challenges to experimentally testing the paradigm of early AD-tau accumulation in AD and its effects.

To address these knowledge gaps, we analyzed Orx^N^ and MCH^N^ in the LHA of human subjects across various stages of the AD-tau continuum. We employed unbiased stereology to examine increases in tau burden and changes in Orx^N^ and MCH^N^ populations. Additionally, we used the nCounter^®^ platform (nanoString^®^ Inc, WA, USA) to profile RNA expression and identify which molecular pathways first exhibit changes in the LHA throughout the AD continuum. This approach aims to elucidate both pathological and compensatory mechanisms associated with AD-tau accumulation in this region, potentially leading to more effective therapeutic strategies for managing sleep-wake disturbances and appetite symptoms in individuals with AD pathology. Our findings indicate that Orx^N^ is the first neuronal population to experience a significant loss in AD, occurring even before neuronal loss in the Enthorynal Cortex (EC) and locus coeruleus (LC).

## Materials and methods

### Participants, selection criteria, and neurological assessment

We carefully selected 69 cases for our study from two cohorts – i) Neurodegenerative Disease Brain Bank (NDBB) at the University of California, San Francisco, and ii) Brazilian BioBank for Aging Studies (BBAS) at the University of São Paulo ^23^. The Institutional Review Boards of both participating institutions approved the study. We adhered to standardized protocols for neuropathological assessments with NIA-AA guidelines to assess the staging of AD neuropathologic changes. All subjects received a comprehensive diagnosis based on the neurofibrillary tangle pathology using the Braak stage scale 0 to 6, established by Braak and Braak ^24^. We also ensured that all cases had complete neuropathological diagnoses ^25^ and available measures of functional cognition based on Clinical Dementia Rating (CDR) **Table 1**.

**Table 1.** Demographic characteristics with the distribution of the cases in each Braak stage, Age of Death, PMI, CERAD score, AD diffused score, CDR score, brain weight, and NIA-Reagan score are represented as mean ± SD.

| Braak Stages | N | Sex | Age of Death (years) | PMI (hours) | CREAD Score | AD Diffused Score | CDR Score | Brain Weight (gram) | NIA-Reagan Score |
| --- | --- | --- | --- | --- | --- | --- | --- | --- | --- |
| BB0 | 8 | 4F, 4M | 61.25 ± 8.4 | 15.64 ± 3.1 | 0.00 ± 0.0**** | 1.00 ± 1.4* | 0.00 ± 0.0* | 1270.50 ± 128.1 | 0.00 ± 0.0*** |
| BB1 | 10 | 5F, 5M | 61.70 ± 7.2 | 19.07 ± 8.2 | 0.25 ± 0.7**** | 0.38 ± 1.1** | 0.14 ± 0.4* | 1219.83 ± 92.1 | 0.20 ± 0.4**** |
| BB2 | 16 | 8F, 8M | 73.75 ± 9.5 | 13.63 ± 5.6 | 0.33 ± 0.5**** | 0.60 ± 1.1** | 0.17 ± 0.4* | 1225.25 ± 103.5 | 0.60 ± 0.5**** |
| BB3 | 4 | 4F, 0M | 72.00 ± 5.4 | 14.38 ± 0.9 | 0.00 ± 0.0*** | 1.00 ± 0.0 | 0.00 ± 0.0 | 1219.33 ± 106.6 | 0.67 ± 0.6*** |
| BB4 | 4 | 2F, 2M | 90.25 ± 2.2 | 09.82 ± 3.2 | 1.75 ± 1.0* | 2.00 ± 1.2 | 0.17 ± 0.3 | 1091.50 ± 37.7 | 2.00 ± 0.0** |
| BB5 | 3 | 2F, 1M | 73.33 ± 8.0 | 11.34 ± 6.1 | 2.67 ± 0.6 | 2.33 ± 0.0 | 0.25 ± 0.4 | 1236.50 ± 39.43 | 3.00 ± 0.0 |
| BB6 | 24 | 10F,14M | 67.67 ± 8.1\$ | 11.85 ± 8.1 | 2.88 ± 0.5 | 2.81 ± 0.5 | 2.36 ± 1.0 | 1093.88 ± 113.8 | 2.92 ± 0.4 |

P-adj values were obtained from a Wilcoxon rank-sum test comparing the demographic characters across different Braak groups where \*p<.05, \*\*p<.01, \*\*\*p<.001 (over BB 6), and \$ p<0.05 (over BB 4). Abbreviations: **BB**, Braak Stage; **CERAD**, Consortium to Establish a Registry for Alzheimer's Disease; **CDR**, clinical dementia rating; **NIA-Reagan**, National Institute on Aging-Reagan Institute criteria for the neuropathological diagnosis of Alzheimer's disease; **PMI**, post-mortem interval.

The demographic details of the cases involved in the study can be found in **Supp. Table S1**. For the study, we included female and male participants aged 50-92 who met specific criteria, including the absence of Lewy body and TDP43, non-AD-related neuropathology insignificant cerebrovascular lesions, and an intact hypothalamus. We excluded individuals with neurological disease, neuropsychiatric diagnosis, or non-degenerative structural pathology. To gather comprehensive data, we did a postmortem stereological evaluation of neural numbers for orexin and melanin-concentrating neurons with CP13 (p-tau) inclusions in the LHA in addition to a probe-based gene expression study. We were mindful of achieving gender balance and maintained a 1:1 female-to-male ratio for our study.

### Tissue processing and immunohistochemistry

Hypothalamic regions containing Orx^N^ and MCH^N^, the area of interest (AOI), were identified using the Allan Human atlas. Formalin-fixed hypothalamic blocks containing the whole AOI were embedded in celloidin ^26,27^ and sectioned serially at 30 µm thickness on the coronal or horizontal plane; section orientation does not affect the optical fractionator probe in stereology. Every 10^th^ tissue section from the AOI was stained with either i) Orexin A and CP13 antibody combo or ii) MCH and CP13 antibody combo and counter-stained with gallocyanin (pH 1.9-2.1) as described previously ^16,17^. In brief, the free-floating serial hypothalamic sections were treated in 0.3% H_2_0_2_ (in methanol) to inactivate endogenous peroxidase and, after that, subjected to 50 mins incubation at 95.7°C in 0.01 M citrate buffer with 0.05% Tween-20 in PBS (pH 6.0) for epitope exposure. To avoid non-specific staining, sections were blocked in blocking buffer (5% milk PBST 0.1% triton X) for 40 mins. Serial sections were double-stained with mouse monoclonal anti-CP13 antibody for phosphor-Ser202 tau (1:1000, a kind gift of Peter Davies, NY, USA) with either rabbit polyclonal anti-Orexin A (1:500, Cat# H-003-30, Phoenix Pharmaceuticals, CA, USA) or rabbit polyclonal anti-MCH serum (1:1000, PBL 234, a kind gift of Joan Vaughan, Salk Institute, CA) overnight at room temperature.

Following overnight primary antibody incubation, the sections were incubated in secondary biotinylated anti-rabbit IgG (1:400, Cat# BA-1100, Vector Labs, CA, USA) and secondary conjugated-HRP anti-mouse IgG (1:400, cat# R-05071-500, Advansta, CA, USA) for 1.5 hours at room temperature. The stained sections were then developed using immPACT DAB Peroxidase (HRP) Substrate Kit (Cat# SK-4150, Vector Labs, CA, USA), Vectastain ABC-AP kit (Cat#AK-5000, Vector Labs, CA, USA), and Vector Red chromogen (Cat# SK-5105, Vector Lab, CA, USA). To estimate the total number of neurons, all sections were counterstained using gallocyanin (Cat# A12936, Alfa Aesar, MA, USA) as a nucleic acid stain at a previously optimized pH (pH 1.9 - 2.1).

We performed double immunohistochemistry of MCH and Orx to rule out possible co-expression of these two markers (**Fig. 1A, B and 2A, B**). **Supp. Fig. S1** depicts representative staining showing the absence of co-expression of these two markers.

**Fig. 1.**
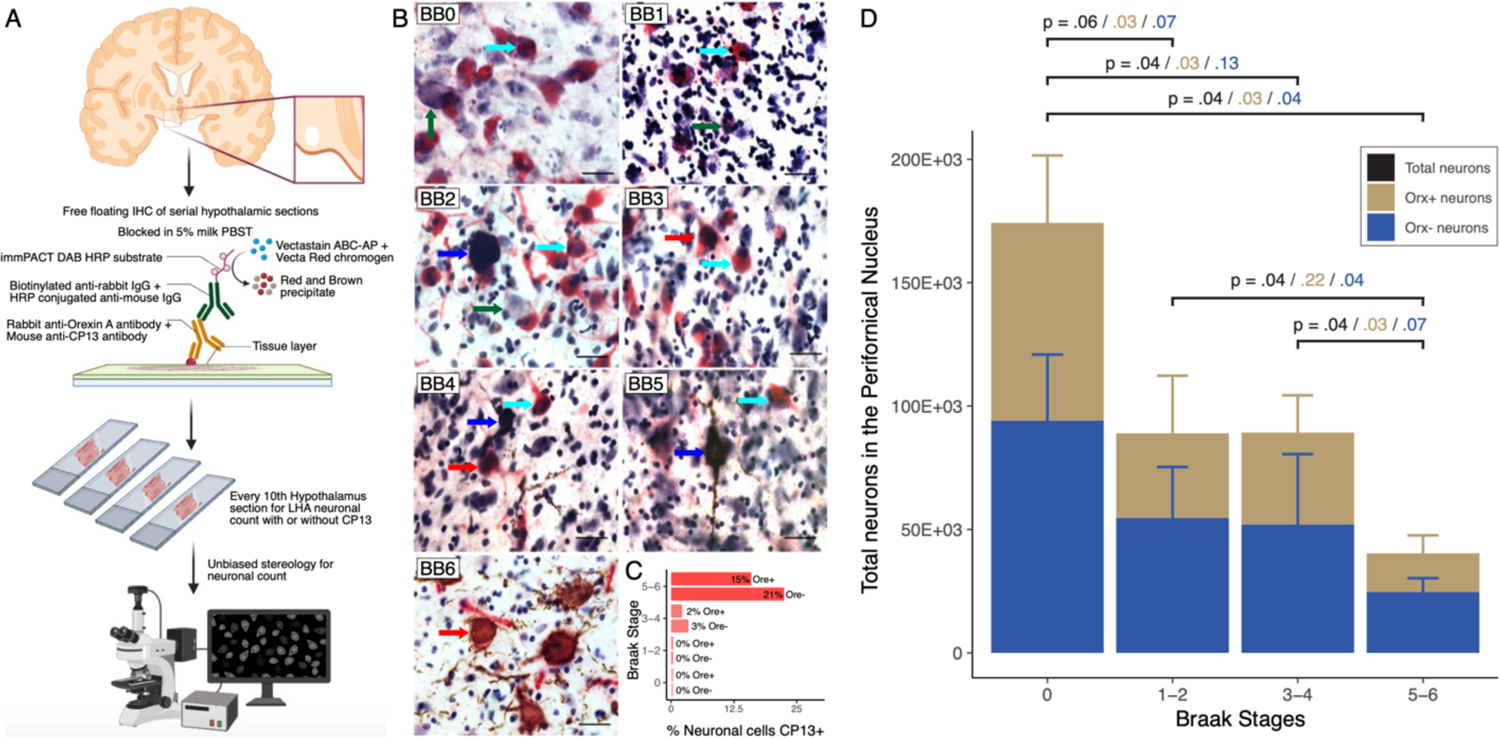
Orexinergic neuronal counts. ***Panel A*** depicts the tissue processing and stereology workflow. ***Panel B*** Representative microphotographs of the perifornical nucleus across Braak groups with Orx-CP13-(**Green arrow)**, Orx-CP13+ (**Blue arrow**), Orx^N^ CP13+ (**Red arrow**), and Orx^N^ CP13-(**Cyan arrow**) neurons at 63x (scale 20µm). The bar plot represents the mean ± standard deviation of the proportion of neurons with Tau inclusion (***Panel C***) and the number of neurons (***Panel D***) in the perifornical nucleus. To determine the significance of the mean difference between the Braak stage of different stereological estimates, we used the Wilcoxon rank-sum test, and p-values were adjusted using the Benjamini & Hochberg test. P-values in black are the comparison between total neuronal count; in light brow are the comparison between ORX^N^, and in blue are the comparison between ORX-.Abbreviations: **CP13+**, pTau (Ser 202) neurons; **CP13-**, pTau (Ser 202) negative neurons; **ORX**^N^, orexin positive neurons; **ORX**-, orexin negative neurons.

### Unbiased stereology

The stereological analyses were performed using the StereoInvestigator v.10 software (MBF Bioscience, VT, USA). Orx^N^ and MCH^N^ were visualized, and live images were captured using a high-resolution camera (MBF, Bioscience, Williston, VT, USA) attached to an Axio Imager.A2 microscope (Carl Zeiss Microscopy, NY, USA). Counting was made for Orx-positive (Orx^N^ Tau-) or MCH-positive (MCH^N^ Tau-), tau-positive (Orx-Tau+ or MCH-Tau+), Orx or MCH and tau-positive (Orx^N^ Tau+ or MCH^N^ Tau+), and double negative neurons (Orx-Tau- or MCH-Tau-) using the optical fractionator probe ^28^. The AOI was delineated at 20x (Plan-APOCHROM 20x/0.8 ∞/0.17, Carl Zeiss Microscopy, NY, USA), and the neuronal counting was performed using a 63x (Plan-APOCHROM 63x/1.4 Oil ∞/0.17, Carl Zeiss Microscopy, NY, USA) objective. The stereological parameters were determined using the “resample–oversample” analysis probes in the StereoInvestigator software ^29^. The guard zone was set at 5 µm, and the dissector height at 12.5 µm; for neuronal count settings, refer to **Supp. Table S2**. The coefficient of error (CE) range was calculated following the methods of Gundersen and Schmitz-Hof ^30,31^.

### Probe-based RNA level expression assay

We utilized the nCounter^®^ digital gene expression assay from nanoString^®^ (nanoString^®^ Inc, WA, USA) to measure changes in gene expression in the LHA across progressive stages of AD. We used the customized human neuropathology panel (PLS_MM_UCSF_1) **Table S3** and glial profile panel to capture and report probes. We followed the manufacturer’s instructions for the gene expression assay by hybridizing 100 ng of total RNA with the codeset overnight at 65°C. The target/probe conjugates underwent automated processing on the nCounter^®^ Prep station, and the unconjugated codesets were removed using nCounter^®^ Master Kit reagents. After purification, conjugated target/probe complexes were immobilized in the nCounter® cartridge for data collection. The nanoString^®^NCTools R software package was utilized to standardize the raw counts and conduct quality checks on data samples collected from each panel used in the study. To gain insights into the overall RNA expression patterns across samples, the run PCA function from the scatter library was employed, facilitating the examination of similarities in expression levels. Principal component analysis (PCA) was then utilized to correlate the primary components (*i.e.*, PC1 and PC2) with covariates and to the visual inspection of technical and biological replicates on the PCA plot **Supp. Fig. S2**. For the differential gene expression analysis, a linear model was employed to compare groups while considering covariates such as Sex, Age of Death, and Binding Density. Notably, Binding Density was included as a covariate strongly correlated with the PCA analysis (or as a highly correlated feature with DV200 and Concentration – **Supp. Fig. S3**). Looking for biological insights, the clusterProfiler library was used to explore gene ontology (GO) and Kyoto Encyclopedia of Genes and Genomes (KEGG) term enrichment through two distinct approaches: over-represented analysis (ORA) and gene set enrichment analysis (GSEA). Furthermore, a detailed examination of gene interactions was conducted using genes identified through enrichment analyses on the StringDB web server.

### Statistical analyses

Mean differences in stereological estimates were assessed between Braak stages for Orx and MCH neurons as population and proportions for each subpopulation (*e.g.*, Orx^N^ Tau-neurons, Orx^N^ Tau+ neurons). Differences in neuronal numbers and proportion were analyzed using the Wilcoxon signed-rank test, with the significance level at .05, and p-values adjusted were obtained using the Benjamini & Hochberg multiple comparison test. All analyses were conducted using R-software (version 4.2; R Foundation for Statistical Computing, Vienna, Austria). In molecular analyses, we deemed covariate correlation significant if Rho exceeded 0.5 and the associated p-value was less than .05. For gene expression, we identified genes as differentially expressed if the p-value, adjusted using Holm correction within the linear model corrected by covariates, was below .05. Similarly, we recognized a term as enriched if its p-value, adjusted by Holm correction, fell below .05.

## Results

### Stereological estimation of orexinergic neurons in the Perifornical nucleus

The optical fractionator method uses systemic random sampling for unbiased neuronal number estimation. The unbiased stereological assessment of the perifornical nucleus (PFN) of the LHA demonstrated a decline in total neuronal number in PFN across the progressive stages of AD. However, the decline reached significance only in the mid-stage of the disease, *i.e.*, in BB 3-4 with a 49% decline (p=.04) over BB 0. By the late stage of the disease (BB 5-6), only 23% of neurons (p=.04) survived in the PFN compared to the BB 0. A simultaneous profound loss of Orx^N^ across the progressive stages of AD, majorly contributing to the global loss of PFN neurons. The decline in Orx^N^ was progressive, starting from BB 1-2 (−58%, p=.03) and continuing to BB 3-4 (−56%, p=.03) and BB 5-6 (−81%, p=.03) over BB 0 **Fig. 1D**.

In conjunction with a decline in Orx^N^, we observed a significant increase in p-tau inclusions in the Orx^N^ and neurons devoid of Orx (Orx-). The proportion of p-tau-inclusion in the Orx-neurons increased in BB 3-4 by 3% and BB 5-6 by 21%. We observed a similar pattern even in the Orx^N^ where 2% of Orx^N^ in BB 3-4 and 15% in BB 5-6 accumulated p-tau **Fig. 1C**. The p-tau inclusion in Orx- and Orx^N^ neurons in BB 3-4 and BB 5-6 was compared with BB 0 and BB 1-2 were significantly higher **Table 2**.

**Table 2.**
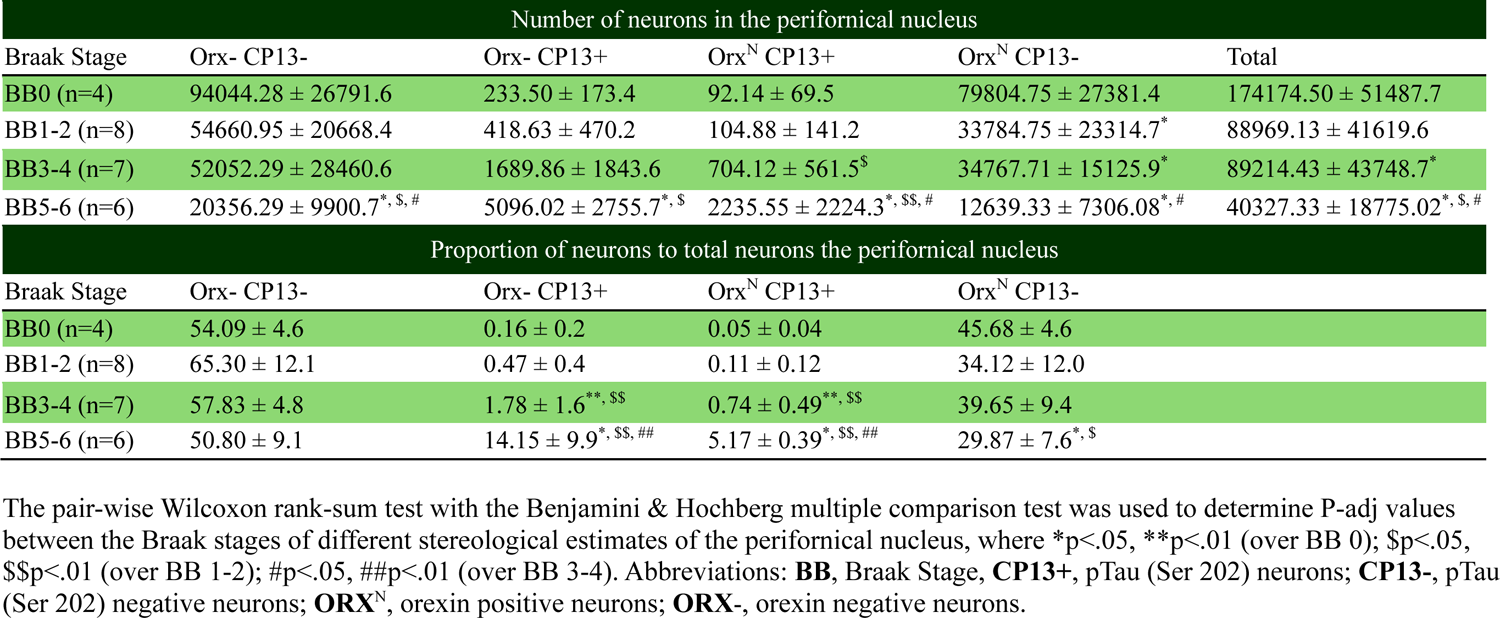
Mean ± standard deviation of stereological estimates of orexinergic neurons in the perifornical nucleus stratified by Braak stages.

### Stereological estimation of melanin-concentrating hormone (MCH) neurons in the LHA

The LHA plays a critical role in sleep-wake modulation through the wake-promoting Orx^N^ and sleep-promoting lateral hypothalamic MCH^N^. Following assessing the Orx^N^, we estimated the total number of neurons in the LHA (LHA^Total^) and MCH^N^. Unlike the significant loss of Orx^N^, the MCH^N^ demonstrated a more preserved profile. The total number of neurons in the LHA demonstrated a nonstatistical significant decline of 25% (p=.1331), while the MCH^N^ population also revealed a 27% (p=.1088) decline in BB 6 over BB 0-2 **Fig. 2D**.

**Fig. 2.**
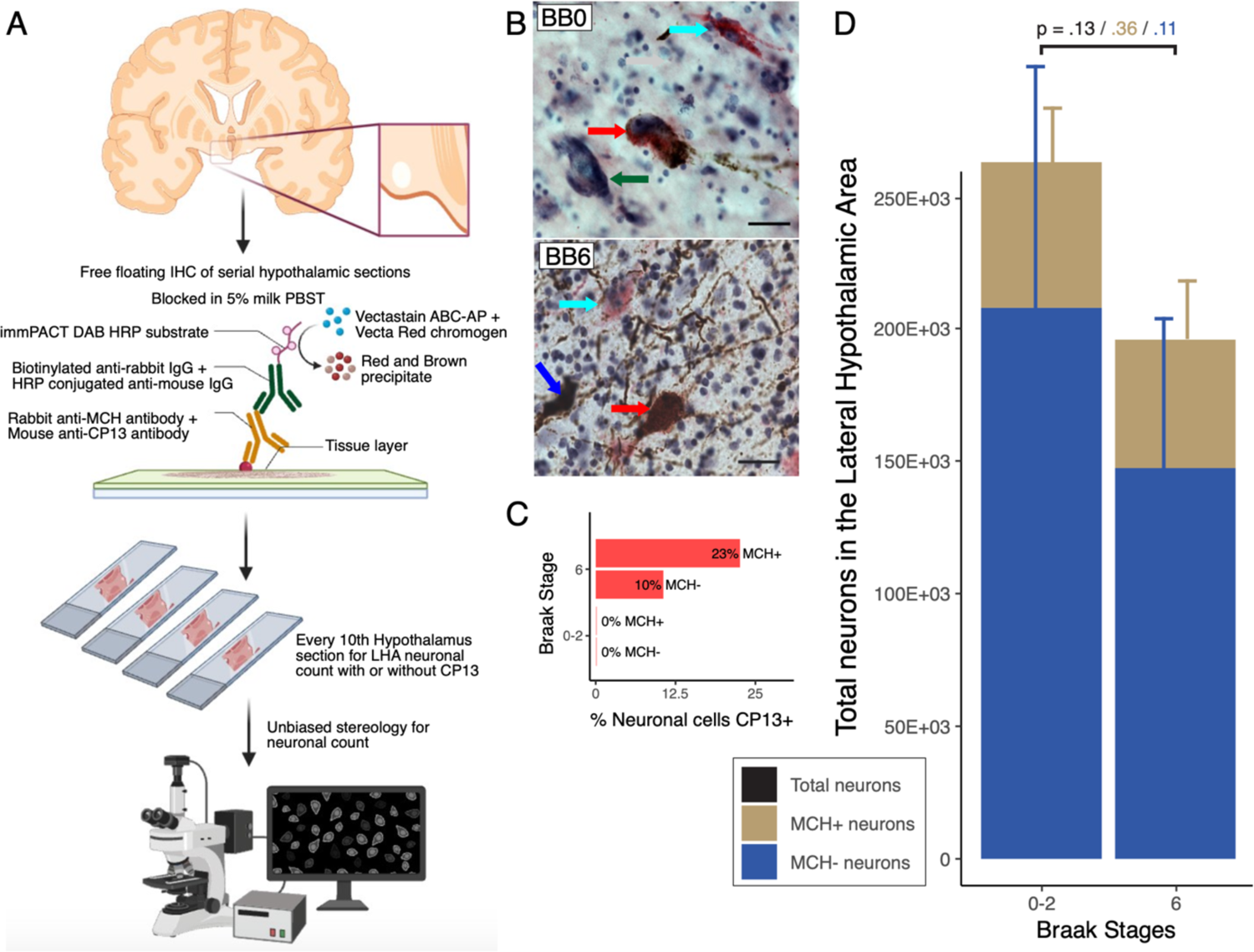
MCHergic neuronal counts. *Panel A* depicts the tissue processing and stereology workflow. *Panel B* Representative microphotographs of the perifornical area across Braak groups 0-2 and 6 with MCH-CP13-(**Green arrow**), MCH-CP13+ (**Blue arrow**), MCH^N^ CP13+ (**Red arrow**), and MCH^N^ CP13-(**Cyan arrow**) at 63x (scale 20µm). Bar plots represent the mean ± standard deviation of the proportion of neurons with tau inclusion (*Panel C*) and MCHergic neurons (*Panel D*) in the LHA. To determine the significance of the mean difference between Braak stage groups of different stereological estimates, we used the Wilcoxon rank-sum test, and p-Values were adjusted using the Benjamini & Hochberg test. P-values in black are the comparison between total neuronal count; in light brow are the comparison between MCH^N^, and in blue are the comparison between MCH-. Abbreviations: **CP13+**, pTau (Ser 202) neurons; **CP13-**, pTau (Ser 202) negative neurons; **MCH**^N^, melanin-concentrating hormone/MCHergic positive neurons; **MCH**-, melanin-concentrating hormone/MCHergic negative neurons.

Further, we counted the p-tau inclusion pattern in the LHA to understand the vulnerability pattern or resilience of MCH^N^ to AD-tau toxicity. We analyzed the proportion of neurons with p-tau inclusion among the MCH^N^ and MCH^-^ neurons. The proportion of p-tau inclusion in MCH^N^ and MCH^-^ demonstrated a significant increase in the late stage of the disease; MCH^N^ demonstrated higher p-tau inclusion (23%) than that of MCH-(10%,) **Fig 3C**. The extent of increase in p-tau inclusion in MCH^N^ (p<.001) was higher than in MCH-neurons (p<.01) **Table 3**.

**Fig. 3.**
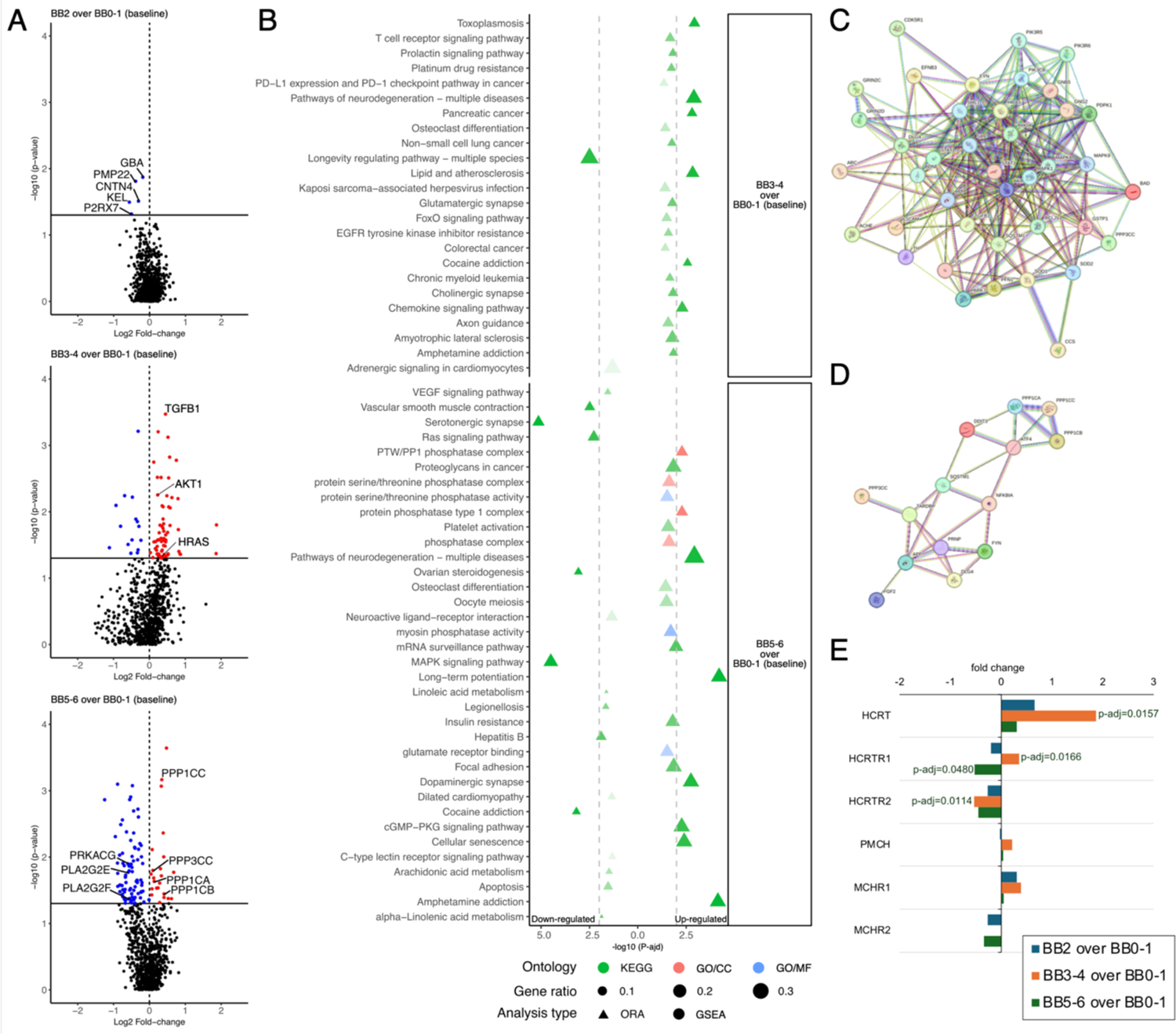
Differentially expressed neuropathology genes in the LHA. ***Panel A*** represents a Volcano plot of differentially expressed genes in Braak stages 2, 3-4, and 5-6 over Braak stages 0-1 (baseline) in the lateral hypothalamic area. The horizontal axis represents the log2FC value; the vertical axis -log10 (p-values); red dots represent up-regulation; and blue dots represent down-regulation. ***Panel B*** Representative functional enrichment analysis (circle, GSEA; and triangle, ORA) of genes with significant GO terms and KEGG pathways associated with the neuropathology panel from Braak stages 2, 3-4, and 5-6 over Braak stages 0-1 (baseline). The color of the bubble indicates the –log_10_ *P*-adj value (p<0.05), and the size of the bubble signifies the gene ratios associated with a term with the molecular function (blue), cellular components (red), and KEGG pathways (Green). ***Panels C*** *and **D*** PPI maps from StringDB using genes associated with enriched pathways. ***Panel E*** Depicts the orexin and MCH-related gene expression with p-values where the vertical axis represents associated genes, and the horizontal axis represents the log2 fold change.

**Table 3.** Mean ± standard deviation of stereological estimates of MCHergic neurons in the lateral hypothalamic area stratified by Braak stage.

| Number of neurons in the lateral hypothalamic area |  |  |  |  |  |
| --- | --- | --- | --- | --- | --- |
| Braak stage | MCH- CP13- | MCH- CP13+ | MCH<sup>N</sup> CP13+ | MCH<sup>N</sup> CP13- | Total |
| BB 0-2 (n=7) | 226288.79 ± 86063.67 | 409.43 ± 484.4 | 14.35 ± 22.26 | 59012.55 ± 18712.41 | 263845.09 ± 111330.28 |
| BB 6 (n=10) | 135762.71 ± 68829.73 | 12835.30 ± 4414.82** | 8245.88 ± 3717.97** | 39822.68 ± 22202.39 | 196666.58 ± 94207.94 |
| Proportion of neurons to total neurons in the lateral hypothalamic area |  |  |  |  |  |
| Braak stage | MCH- CP13- | MCH- CP13+ | MCH<sup>N</sup> CP13+ | MCH<sup>N</sup> CP13- |  |
| BB 0-2 (n=7) | 78.64 ± 2.19 | 0.15 ± 0.18 | 0.004 ± 0.006 | 21.21 ± 2.11 | |
| BB 6 (n=10) | 67.30 ± 5.6** | 7.68 ± 3.14** | 5.91 ± 6.6** | 19.12 ± 5.92 | |

The Wilcoxon rank-sum test with the Benjamini & Hochberg multiple comparison test was used to determine P-adj values between the Braak stages of different stereological estimates of the LHA, where \*\*p<0.01 (over BB 0-2). Abbreviations: **BB**, Braak Stage, **CP13+**, pTau (Ser 202) neurons; **CP13-**, pTau (Ser 202) negative neurons; **MCH<sup>N</sup>**, melanin-concentrating hormone/MCHergic positive neurons; **MCH-**, melanin-concentrating hormone/MCHergic negative neurons.

### Identification of differentially expressed genes in the LHA

To investigate the alterations in expression patterns in the LHA associated with neuronal vulnerability identified in stereological assays, we utilized the nCounter^®^ platform (nanoString^®^ Inc, WA, USA) throughout different stages of AD. Initially, we employed a customized Neuropathology panel (see methods). On this customized Neuropathology panel (composed of 790 genes), we observed a progressive increase in the number of differentially expressed genes (DEGs), with 5 DEGs identified in BB2, 80 DEGs in BB 3-4, and 105 DEGs were found in BB 5-6, compared to BB 0-1 **Fig. 3A**. No significant enrichment in GO/KEGG pathways was detected for the 5 DEGs in BB 2, all of which PMP22, CNTN4, P2RX7, KEL, and GBA were downregulated. However, BB 3-4 and BB 5-6 exhibited various up- and down-regulated pathways **Fig. 3B**. In BB 3-4, genes associated **Fig. 3C** with TGFβ signaling, HRAS, and AKT1, which are implicated in microglial changes ^32^, amyloid plaque-associated dendritic spine loss in AD mouse models ^33^ and phosphorylation that modulates various process in development and progression of AD ^34^, respectively, enriched pathways such as pancreatic cancer, chronic myeloid leukemia, FoxO signaling pathway, osteoclast differentiation, colorectal cancer, neurodegeneration, cholinergic synapse, chemokine signaling pathways, amyotrophic lateral sclerosis, and glutamatergic synapse. The observed glia’s chances in BB 3-4 seem to be started in BB 2 with PMP22 and GBA genes, which are recognized to participate in the myelination process ^35^. In BB 5-6, up-regulation of the protein phosphatase family (*i.e.*, PPP1CB, PPP1CC, PPP1CA, and PPP3CC) led to the enrichment of pathways related to various phosphatase processes and the dopaminergic synapse. Furthermore, in BB 5-6, a gene cluster containing PRKACG, PLA2G2F, and PLA2G2E **Fig. 3D** enriched, among others, the MAPK signaling pathway, a well-known pathway associated with neuronal death in AD ^36^. Examining specific genes associated with neuropeptide transmission in LHA Orx^N^ and MCH^N^ (*i.e.*, HCRT and PMCH genes) along with neuropeptide receptors (*i.e.*, HCRTR1, HCRTR2, MCHR1, and MCHR2 genes), alterations in the expression of Orx-related genes were evident **Fig. 3E**. Minor fluctuations in MCH transmitters and receptors did not attain statistical significance. Intriguingly, a notable upregulation of HCRT expression was observed in BB 3-4 compared to BB 0-1, despite the noted selective vulnerability in the stereological data.

Upon recognizing the direction of molecular changes toward glial adaptation and the limitation of the Neuropathology panel to detect genes that could explain glial changes, we applied the nanoString^®^ Glia Profiling Panel to the same samples. As expected, we observed a substantial number of DEGs, even in BB 2 (n = 78), within 784 targeted genes of the glia panel, compared to no change in the neuropathology panel at the same BB **Fig. 4A**. Almost all enriched GO/KEGG terms **Fig. 4C** in the BB 2 were influenced by the downregulation of ATP1B1, ATP2A2, ATP5MC3, ATP6V0D1, ATP6V1A, ATP6V1E1, ATP6V1G2, and ATP6V1H, predominantly associated with the mitochondrial dysfunction, oxidative phosphorylation process, lysosomal dysfunction, and innate immune modulation ^37,38^ **Fig. 4D**. Mitochondrial dysfunction and oxidative stress are subjects of ongoing discussion in AD research ^39–41^. The Graft-versus-host disease pathway is another enriched pathway in BB 2 that calls our attention, enriched by the genes KIR2DL3, KIR3DL1, and GZMB. Killer cell immunoglobulin-like receptors (KIRs) are transmembrane glycoproteins expressed by natural killer cells and certain T cell subsets.

**Fig. 4.**
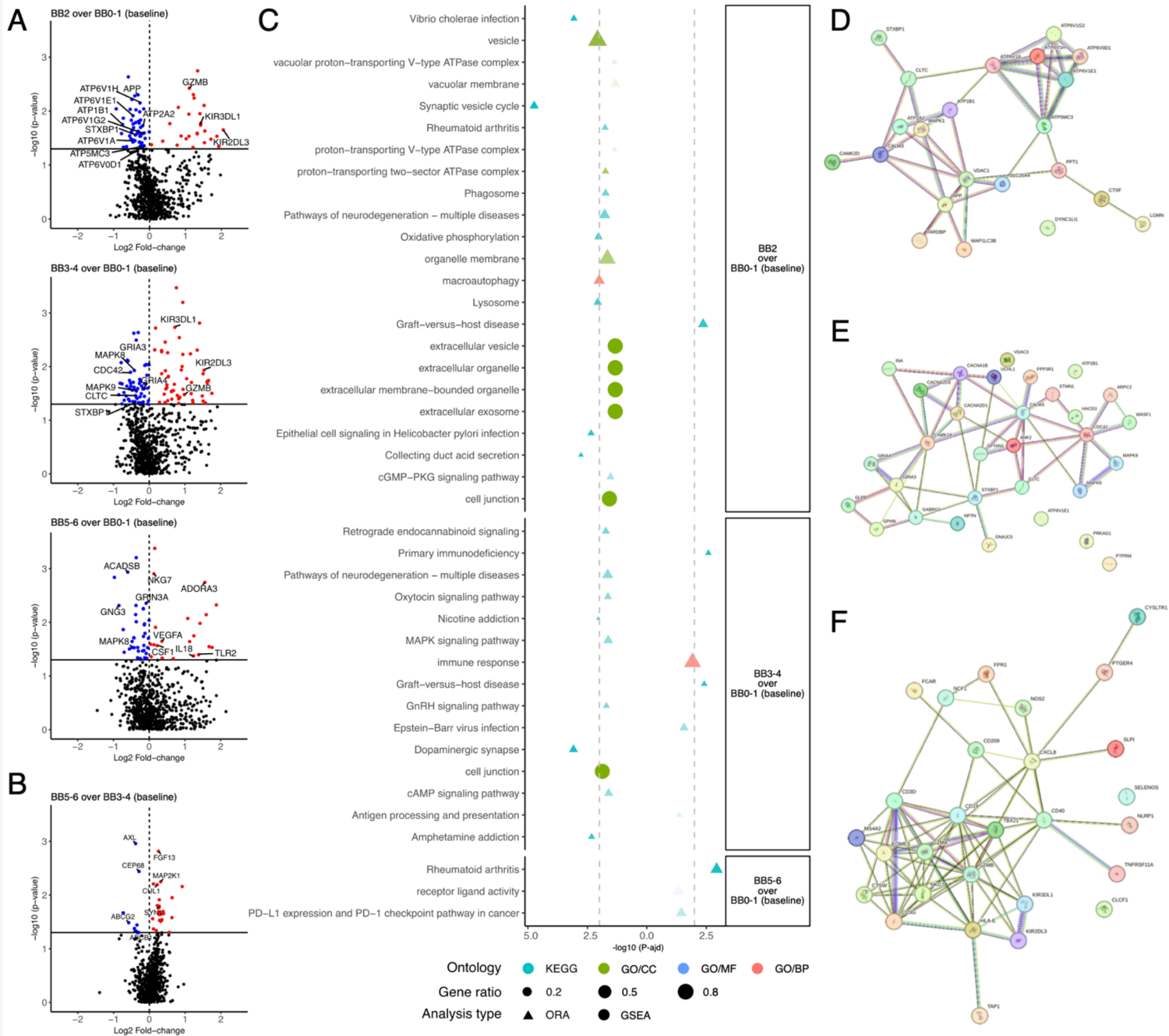
Differentially expressed glial genes in the lateral hypothalamic area. ***Panel A*** represents a Volcano plot of differentially expressed genes (DEGs) in Braak stages 2, 3-4, and 5-6 over Braak stages 0-1 (baseline) in the lateral hypothalamic area; and ***Panel B*** represents a Volcano plot of DEGs in Braak stage 5-6 over Braak stages 3-4. The horizontal axis represents the log2FC value; the vertical axis -log10 (p-values); red dots represent up-regulation; and blue dots represent down-regulation. ***Panel C*** Representative functional enrichment analysis (circle, GSEA; and triangle, ORA) of genes with significant GO terms and KEGG pathways associated with the glia panel from Braak stages 2, 3-4, and 5-6 over Braak stages 0-1 (baseline). The color of the bubble indicates the –log_10_ *P*-adj value (p<0.05), and the size of the bubble signifies the gene ratios associated with a term with the molecular function (blue), cellular components (red), and KEGG pathways (Green). ***Panels D****, **E**, and **F*** PPI maps from StringDB using genes associated with enriched pathways.

Following the progression of the disease, BB 3-4 exhibited the highest number of DEGs for this panel (n = 107), with a clear distinction between down-regulated genes associated with MAPK signaling and the dopaminergic synapse **Fig. 4E**, and up-regulated genes linked to Graft-versus-host disease and primary immunodeficiency pathways **Fig. 4F**. While in BB 5-6, a total of 57 genes displayed altered expression compared to BB0-1. Within these alterations, genes CSF1, IL18, TLR2, and VEGFA were implicated in the Rheumatoid arthritis pathway, indicating an inflammatory response alteration. Notably, a significant late change in BB 5-6, when compared with BB 3-4, was the downregulation of terms associated with ABC transporters, previously identified as key players in AD ^42^, and the up-regulation of terms linked to cAMP signaling, a messenger system potentially implicated in AD-related neurodegeneration ^43^ **Fig. 4B**. The high number of DEGs in the BB 2 supports the hypothesis that microglial adaptation and an immune response preceding neuronal death commence in the initial stages.

To ascertain whether the alterations observed in BB 5-6 of typical AD and atypical AD exhibit similarities, we categorized BB 5-6 cases and conducted tests against BB 0-1. Within the Neuropathology panel, we noted the upregulation of pathways associated with the protein phosphatase family (*i.e.*, PPP1CB, PPP1CC, PPP1CA, and PPP3CC genes) in both AD phenotypes. However, in atypical AD cases, these genes were accompanied by numerous other genes from the same pathway, suggesting a heightened enrichment of terms related to phosphatase processes and the dopaminergic synapse. The second gene cluster previously mentioned, which was downregulated in late AD and related to the MAPK signaling pathway (such as PRKACG, PLA2G2F, and PLA2G2E), was exclusively observed in atypical cases.

Additionally, in atypical AD cases, BB 5-6, compared to BB 0-1, revealed several genes following the CD40 gene in the enrichment of pathways related to microglial activation. Conversely, within the glia panel, the VEGFA gene, previously implicated in the enrichment of the Rheumatoid arthritis pathway and indicative of an alteration in the inflammatory response, remained present only in typical AD cases pathways, alongside several other neuroprotective promoters like EGF and IGF1 genes. Each phenotype takes divergent routes to induce neuroinflammation, leading to subsequent neurodegeneration. This observation underscores the complexity of the molecular mechanisms underlying AD and highlights the need for further investigation into the specific pathways involved in each phenotype.

## Discussion

The present study builds upon our previous findings of profound loss of wake-promoting neurons in the subcortical areas in AD ^16^. We systematically studied two distinct neuronal populations in the LHA that play a critical role in regulating sleep and wakefulness. Our findings show a decline in the LHA neuronal population starting as early as BB 1-2. This decline is more pronounced in Orx^N^ compared to the neighboring Orx-, MCH- and MCH^N^. Unbiased stereology revealed the pattern of Orx^N^ decline during Alzheimer’s disease progression, revealing a significant neuronal loss in the Orx^N^ population at BB 1-2 and 5-6. Meanwhile, in BB 3-4, neuronal loss plateaued. Ultimately, we found that approximately 20% of Orx^N^ survived in the PFN of LHA. The MCH^N^ presented a contrasting trend with remarkable resistance to tau-toxicity characteristic of Alzheimer’s disease.

The LHA plays a critical role in maintaining the sleep- and wake cycle; approximately 25-44% of AD patients suffer from sleep-wake disturbance ^44,45^. Excessive daytime sleepiness, reduction in sleep efficiency with fragmented sleep at night, a decrease in the percentage of time of NREM3 stages and REM sleep, and an increase in wake after sleep onset, indicate that the wake-promoting system gets severely affected in AD ^17,18^. The Orx^N^ along with the noradrenergic neurons of the LC and histaminergic neurons of the tuberomammillary nucleus (TMN), maintain wakefulness ^4–6^. While the MCH is important in maintaining REM sleep ^46,47^. Loss or destruction of MCH neurons promotes arousal and wakefulness, with a decrease in non-REM sleep.

Hyperactivity of MCH neurons can lead to more REM sleep at the expense of slow wave sleep (SWS) ^48,49^. Although the MCH^N^ were preserved in the LHA in AD, we found a substantial increase in p-tau accumulation in parallel with Orx^N^.

Aging, per se, is considered the primary risk factor for neurodegenerative disorders, including AD ^50^. Using male Fisher 344/Brown Norway F1 hybrid rats, Kessler *et al.* demonstrated that age-dependent neuronal loss in the LHA was not global. They quantified the immunoreactivity of Orx^N^ and MCH^N^ in the male rats and demonstrated a significant decline (*i.e.*, ∼40%) in Orx^N^ immunoreactivity to the medial and lateral areas of the fornix. Meanwhile, the decline in MCH^N^ immunoreactivity was restricted to the area medial to the fornix ^51^. In parallel, an age-dependent significant decline (p=.023) was evident in Orx^N^ in older adults (*i.e.*, 48-60 years) over young adults (*i.e.*, 22-32 years) in a cohort of normal aging in humans without any neuropathology ^52^.

Paradoxically, an increase in Orx levels in CSF (Orx^csf^) in moderate to severe AD patients with sleep impairment and cognitive decline has been reported in recent studies. This elevated Orx^csf^ correlates positively with CSF Aβ42, t-tau, and p-tau levels ^18,20–22^. Moreover, the extent of cognitive impairment was more pronounced in AD patients with various neuropsychiatric with elevated Orx^csf^ levels ^15,20^. In parallel with increased Orx^csf^ levels, plasma Orx levels also demonstrated elevated levels in AD patients ^53^. Pharmacological manipulation of the orexinergic neurotransmitter system across various animal models has demonstrated neuroprotective potential. Either administration of Orx or manipulation of Orx receptors has been shown to stabilize glucose metabolism, reduce neuroinflammation and oxidative stress ^54–56^, and mediate neuroprotection by activating ERK1/2 and AKT pathways ^54,57^. In addition to sleep regulation, MCH^N^ also regulates appetite and hippocampus-dependent memories ^58^. A study on the rodent model demonstrated that REM sleep-dependent inhibition of MCH neurons impaired hippocampal-dependent memory ^58^. Further, Calafate *et al.* demonstrated that MCH neuronal loss is an early event in AD, downregulates synaptic transmission, modulates firing rates in the hippocampal neurons, and induces sleep defects in the APP^NL-G-F^ mouse model of AD ^59^. Despite extensive studies in experimental animals, the knowledge gap for MCH neurons in humans across various neurodegenerative diseases is substantial.

AD-associated genes, including TREM2 and APOE, are highly expressed within the innate immune system, and their pivotal function lies in modulating the neuroinflammatory reaction of microglia to tau pathology ^60^. Molecular alterations identified the downregulation of v-ATPase, Na^+^/K^+^ATPase, Ca^+^ATPase, and mitochondrial ATPase gene in BB 2. Allosteric activation of ATPV0D1, a key component of v-ATPase, is critical for lysosomal acidification. ATP6V0D1 knockdown leads to a significant decline in cell survival and loss of lysosomal acidity in PC12 cells and induces neurodegeneration in rodent models of diabetic neuropathy ^61^. In addition, inhibition of ATP6V0D1 can inhibit macrophage autophagy. A recent study demonstrated inhibition of ATPV0D1 expression via virulence-associated protein A (VapA) inhibits macrophage autophagy in J744A.1 cell lines ^62^. Dysfunctional microglial autophagy is linked to neuroinflammation and contributes to the pathogenesis of AD ^63,64^. Our results indicated that microglial changes reflected an early impact on the LHA in AD and the selective vulnerability of Orx^N^. Furthermore, microglia may actively contribute to synaptic dysfunction by aberrantly phagocytosing synaptic components of neurons affected by tau pathology ^65^. The Na^+^/K^+^ATPase downregulation was seen in CSF of AD patients ^66^. Na^+^/K^+^ATPase plays a crucial role in maintaining transmembrane ion gradient and neuronal excitability. Studies have suggested that impairment in Na^+^/K^+^ATPase activity is associated with oxidative stress and Amyloid-β and contributes to AD pathology ^67,68^. The orexinergic system’s extensive projections to various brain regions, including noradrenergic neurons of the LC, histaminergic neurons of the TMN, serotonergic neurons of the Raphe Nuclei, and dopaminergic neurons of the Ventral Tegmental Area, as well as its innervation of cholinergic and noncholinergic neurons in the basal forebrain and cortex, highlight its regulatory role in multiple neurotransmitter systems ^69^. Synapse pathways related to Glutamatergic, Cholinergic, Serotonergic, and Dopaminergic had changes observed in the molecular data in BB 3-4 and BB 5-6 compared to BB 0-1. For example, the upregulation of HCRT during AD progression could suggest a compensatory mechanism to sustain this orexinergic system. In the same way, the downregulation of HCRTR1 and HCRTR2, possibly linked to neighboring MCH^N^, may indicate an excessive neurotransmitter supply. This Orx^N^ hyperexcitability at BB 3-4 could also be identified by the significant decline in ATP1B1 expression observed. A decline in ATP1B1 expression may lead to hyperexcitability and an excitotoxic cellular response-mediated neurodegeneration ^67,70^. Increased Orx levels have also been associated with tau hyperphosphorylation (T231) in primary hippocampal pyramidal neurons, which HCRTR1 inhibitor SB334867 reverted ^53^. Further investigations are warranted to ascertain whether Orxinergic neuronal vulnerability arises solely from pathological stress or from an augmented production to maintain the overall orexinergic system in the brain.

In conclusion, the Orx^N^ of the LHA demonstrated a selective neuronal vulnerability to AD-specific p-tau and is the first neuronal population to die preceding the loss of LC neurons in AD. At the same time, the neighboring MCH neurons were comparatively more resilient, with a substantial p-tau burden. Molecular data alongside the stereological analysis indicates glial profile change or neuroinflammation as the initial driver of the observed Orx^N^ loss in the neuromodulatory subcortical LHA. On the other hand, the second decline in Orx^N^ after BB 3-4 was associated with the loss of VEGF pathways, a decline in neurotransmission patterns with increased neuroinflammation, and possibly due to hyperexcitability of Orx^N^. Interventions preventing Orx^N^ loss and inhibiting p-tau accumulation in the LHA could prevent neuronal loss in AD and, perhaps, the progression of the disease.

## Acknowledgment

This work was supported by grants from Tau Consortium/Rainwater Charity Foundation, NIA R01AG060477, NIA R01 AG064314, NIA K24 AG053435 (Grinberg).

## Conflict of Interest

The authors do not have any conflict of interest to disclose.

## Notes

### Competing Interest Statement

The authors have declared no competing interest.

### Summary of Updates

Rectification of typographical errors and type settings. Parenthesis was removed from around the fig and legends in the text. Revised the supp. Table 2.

