## Supplementary figures for "The wake- and sleep-modulating neurons of the lateral hypothalamic area demonstrate a differential pattern of degeneration in Alzheimer’s disease"

**Supp. Figures:**

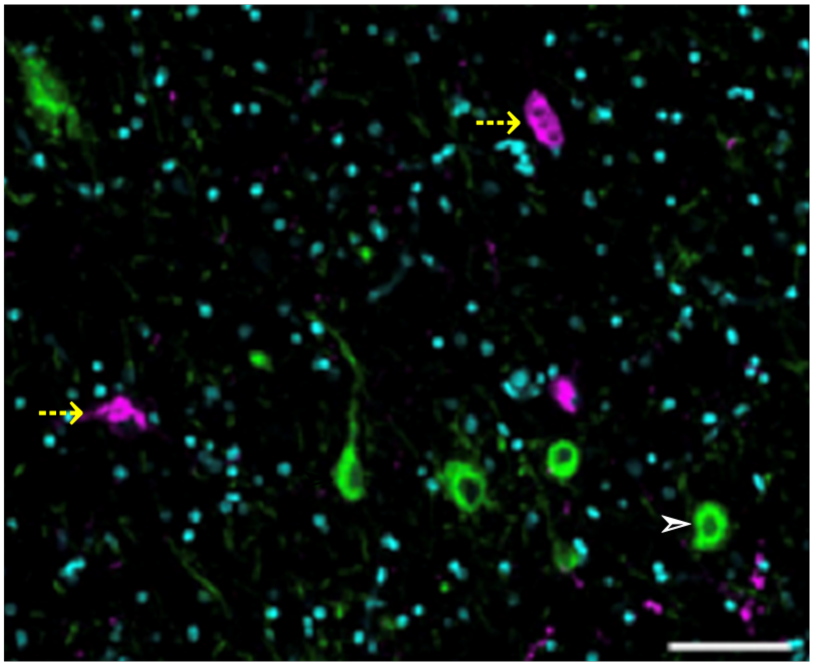

**Supp. Fig 1** Neurons in the lateral hypothalamic area (LHA) do not co-express MCH and orexin. The figure represents double-staining immunohistochemistry for MCH (white arrowhead) and orexin (yellow dotted arrow) in the LHA of a control subject at 20x (scale bar 100um).

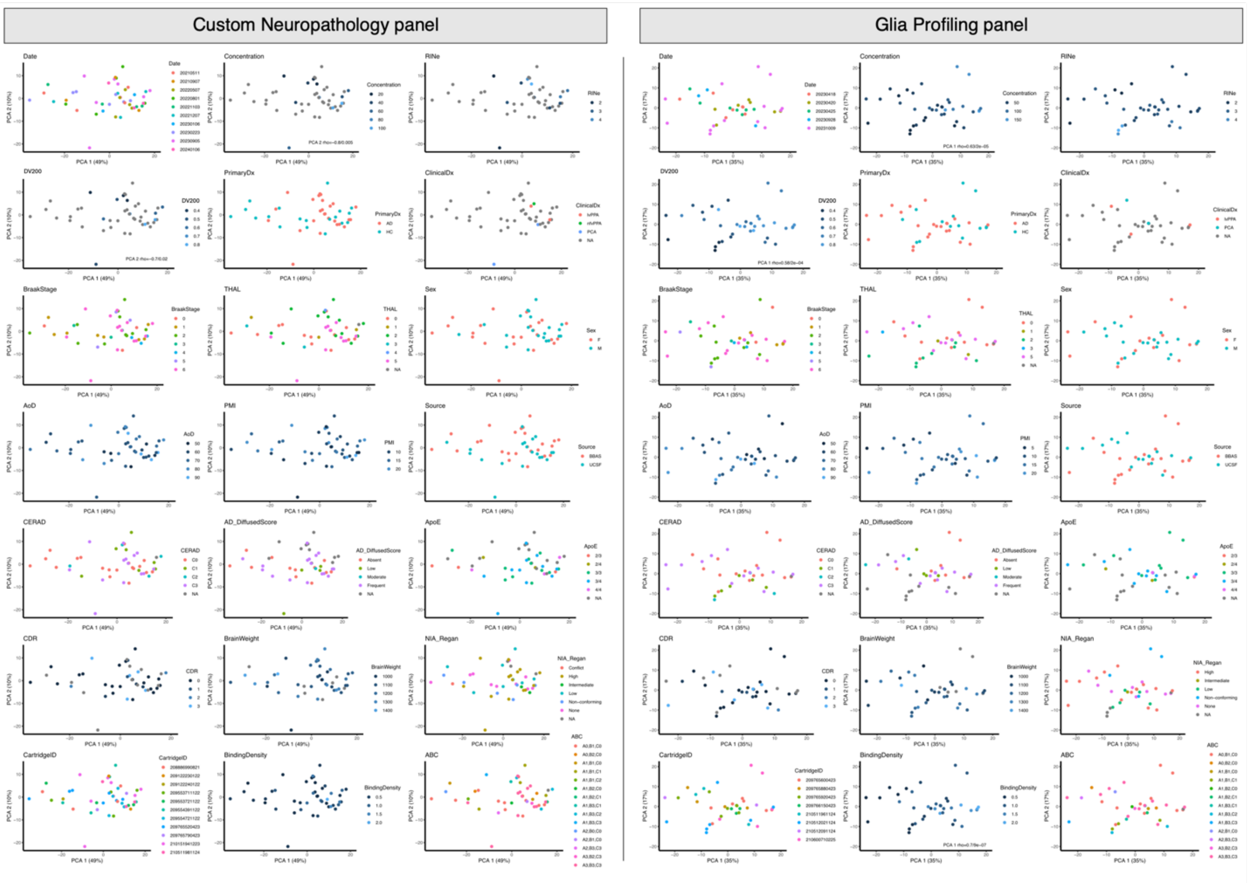


**Supp. Fig 2.** Principal component analysis and correlation analysis of the primary components (*i.e.*, PC1 and PC2) with covariates.

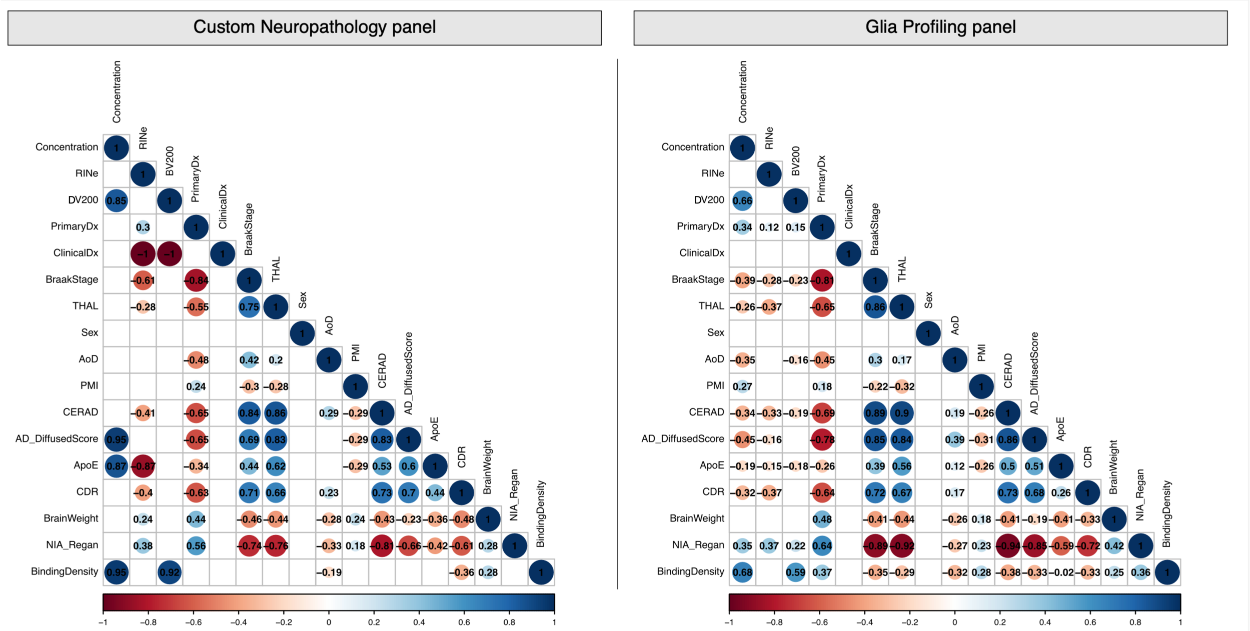


**Supp. Fig 3.** Correlation analysis of covariates. Correlation analysis of covariates revealed that binding density had a strong correlation with PC1 in the Glia Profiling Panel. Additionally, DNA concentration and DV200 showed a strong correlation with PC2 in the Neuropathology Panel. Since binding density is also strongly correlated with DNA concentration and DV200, it was used as a correction factor in the linear model to identify differentially expressed genes.
